# Time-Periodically Driven Brownian Motion of Rigid Rod in one dimensional space

**DOI:** 10.1101/297861

**Authors:** M. A. Shahzad

## Abstract

In this paper we discuss a simple theoretical approach, taken from the theory of stochastic processes to understand the basic phenomenology of protein translocation through a flickering pore. In this theoretical approach we investigate the dynamics of Brownian particle driven by a periodically driving force. This toy model is further extended by considering the Langevin equation with constants drift and time dependent variance. Using the first passage time theory we derived the formalism for probability density function to comprehend the translocation process occurring in the presence of fluctuating environment.

---

Translocation of biomolecule through nanoscale pores is a complex physical process. Study of translocation process is importance for various biological phenomena such as injection of viral DNA, RNA transfer across nuclear pores, and protein transport through membrane nanopores. Translocation process also play an important role to understand the DNA sequencing and sorting [1–10].

Recently, some theoretical and experimental research groups vary the width of the pore during the translocation process. In such translocation process, the dynamical nature of the pore enable the polymer chain to translocate more efficiently as compare to the translocation of polymer in static pore [11–15]. On the experimental side, it has been shown that the translocation of DNA through a nanochannel can be modulated by dynamically changing the cross section of an elastomeric nanochannel device by applying mechanical stress [16–20]. Time dependent driving force are also used in the translocation process [21–25]. There are some biological examples of such fluctuating environment in translocation are the nuclear pore complex, which plays an essential role in nucleocytoplasmic transport in eukaryotes [26]. Moreover, using an alternating electric field (time-dependent driving force) in the nanopore has been implied as a source for DNA sequencing [28]. P. Fanzio, et al., [17] has shown that the DNA translocation process through an elastomeric nanochannel device can be altered by dynamically changing its cross section. The opportunity to dynamically control the nano-channel dimension open up new possibilities to understand the interactions between bio-molecules and nano-channels, such as the dependence of the translocation dynamics on the channel size, and the effects of moving walls [11]. Here we discuss a theoretical model to understand the translocation processes through a vibrating nanopore.

We consider a Brownian rigid rod particle of length *λ* on one space dimension, driven by a time-periodic driving force, *F* (*t*) = *F* cos(*ωt* + *θ*), where *F* is the amplitude of force, *ω* is its frequency and *θ* is the phase shift. The translocation of such rigid rod is illustrated in Fig. (1). Assuming that the rigid rod moves as a free Brownian particle in space (*U(x)* = 0) [5], the Langevin equation for large friction is given by

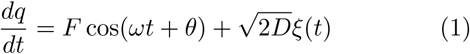

where *ξ*(*t*) is a white noise force that has zero mean, 〈*ξ*(*t*)〉 = 0. The white noise force *ξ*(*t*) is delta correlated, 〈*ξ*(*t*)*ξ*(*t^′^*)〉 = *Dδ*(*t – t^′^*)〉, where *D* is the diffusion coefficient.

In the limit of low diffusion *D* → 0, the above equation becomes deterministic, that is

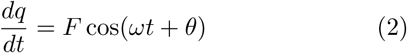

**FIG. 1.**
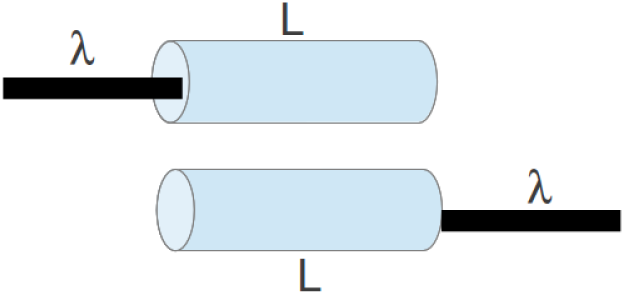
Translocation of rigid rod particle of length *λ* through a cylinder having length *L*.

Using the initial condition at the edge of channel, *q*(0) = 0 and *q*(*τ*) = *L*, the integral form of the above equation is

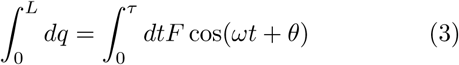

Carrying out the integration, one can easily obtain the formula for the first passage time

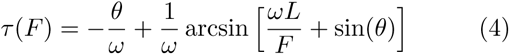

Figure (2) shows the plot of the translocation time *τ* (*F*) as a function of constant force *F* obtained from the above equation. When the force is low enough (*F <* 10), there is an abrupt change in time, the time *τ* increases with the decrease of the constant force *F*. However, the time becomes insensitive with respect to force change in the limit of *F* large.

We consider the encounter of the rod-like particle at the pore and its eventual escape into either the cis or trans side as a one-dimensional stochastic process of the particle under the action of applied driven force *F* across the channel. The equation of motion for the state variable *q*(*x*) can be formulated in the form of a stochastic differential equation (Langevin equation) given by

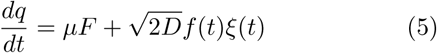

**FIG. 2.**
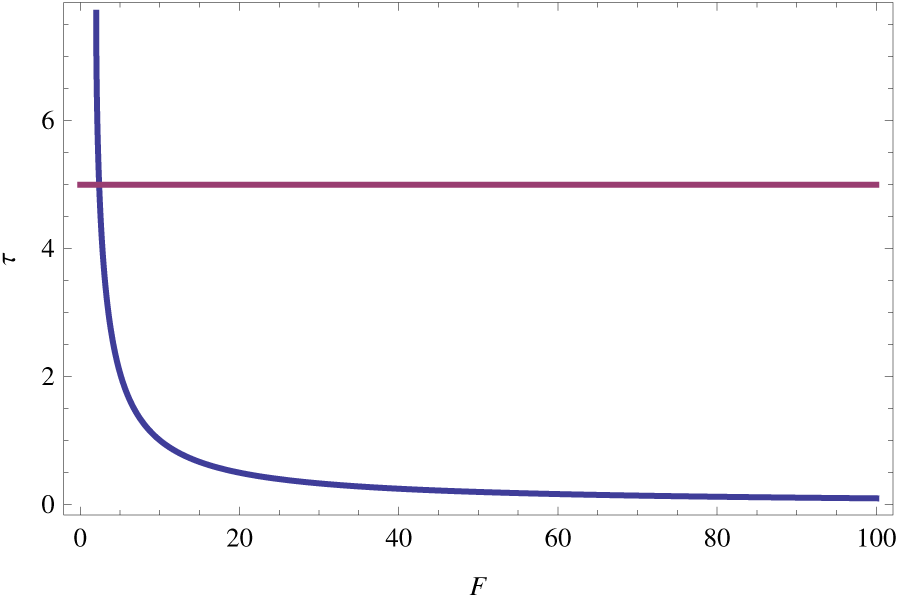
Translocation time as a function of force *F* across a channel. The translocation time decrease exponentially with the increase of constant force.

where *F* is constant force, the friction *µ* is referred to as the drift coefficient, while *D* is called the diffusion coefficient, and *ξ*(*t*) is a white noise. The fluctuating variance *σ*(*t*)^2^ = *f* (*t*) = *A* cos(*ωt* + *θ*), where *A* is amplitude, *ω* is the frequency of the fluctuating variance. The schematic diagram of the such process are shown in Fig. (3).

**FIG. 3.**
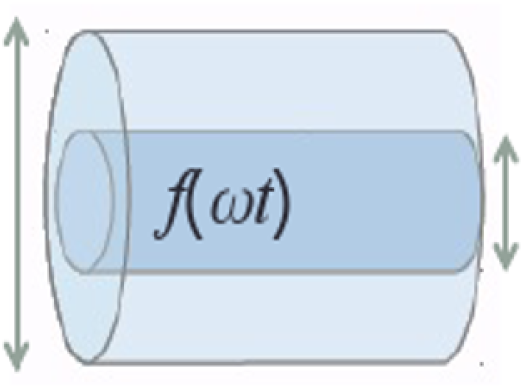
Fluctuating rigid rod particle of length *λ* around the minima.

Using the initial condition *q*(0) and the boundary passage time can be easily obtained by using Eq. (3.26),

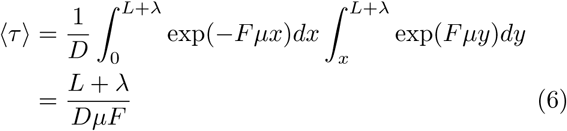

The simplest time-discrete approximation of the Langevin equation is the stochastic generalization of the Euler approximation

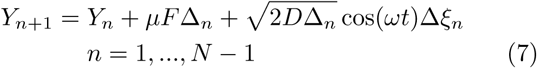

with the initial value *Y*_0_ = *x*_0_. The time step are ∆_*n*_= *t*_*n+1*_ – *t*_*n*_, and ∆ *ξ*_*n*_ = *ξ*(*t*_*n+1*_) – *ξ*(*t*_*i*_).

In Fig. (4) we plot the first passage density obtained via direct simulation of Eq. (5). The distribution of the first passage density shows many peaks which are locked to the variance modulation (green dotted line in Fig. (4)). The Langevin equation with time dependent diffusion term *σ*(*t*) is given by

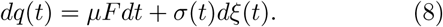

**FIG. 4.**
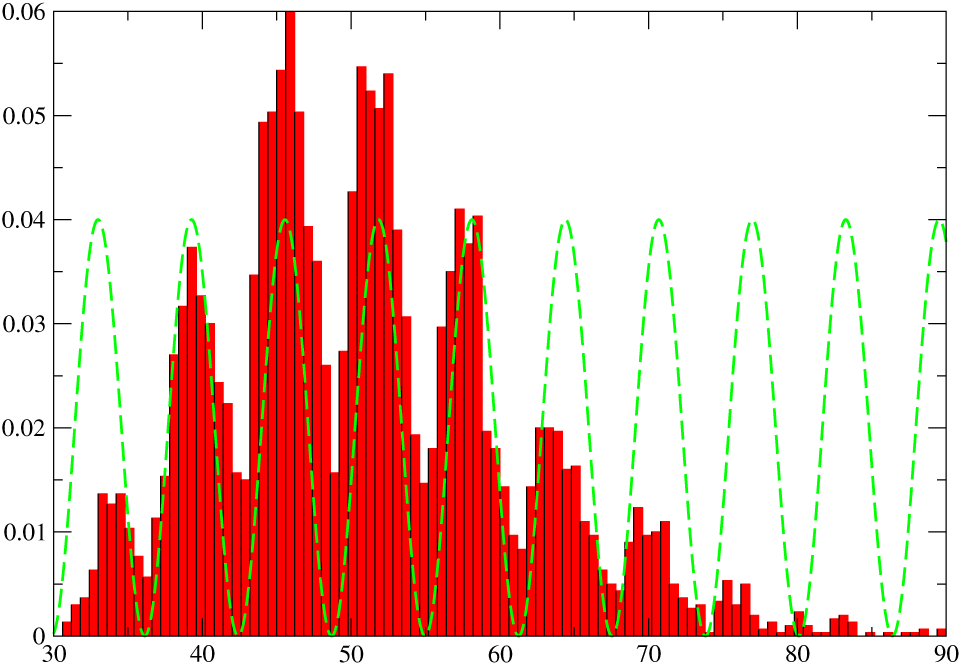
The distribution of the first passage time at *ω* = 5, *θ* = 0 and *F* = 1, computed via numerical simulation of Eq.(5). The distribution displays many peaks locked to the variance modulation, *f* (*ωt*) = *A* cos(*ωt* + *θ*).

where *q* is reaction coordinate. A reaction coordinate *q* defined via the piecewise function [34]

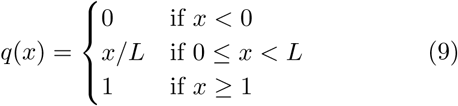

where *x* is the coordinate of the particle. Here, *q* = 0 when particle is on the cis-side of the channel, *q* = 1 when the particle is completely on the trans-side, and 0 *< q <* 1 if there is any particle inside the channel. The expression for collective coordinate becomes

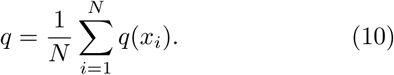

The associated Fokker-Planck equation [31, 32] describing the evolution of the probability density function (PdF) of *q*(*t*) can be expressed as

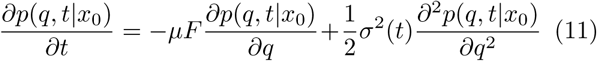

where *p*(*q, t x*_0_) is transition PdF with initial condition *P* (*q,* 0) = *δ*(*q*). Changing the original variable *q* and *t* into

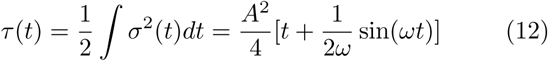

with *σ*(*t*) = *A* cos(*ωt*), and

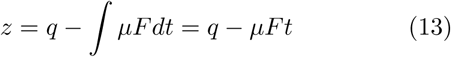

the normalized solution of Fokker Planck equation in the unbounded space becomes,

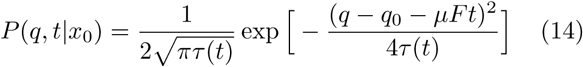

We used the method of images for solving the boundary value problems with time dependent diffusion coefficient. To solve the FP equation using the method of images, we place a mirror source at q=2 (as shown in Fig. (5)) such that the solution of the FP equation emanating from the original and mirror sources exactly compensate each other at the position of the mirror source at each instant of time. This implies that the initial condition *P* (*x,* 0) = *δ*(*x*) must be modified to

**FIG. 5.**
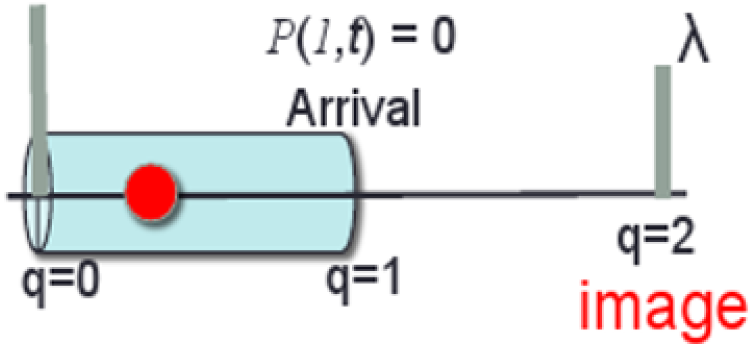
Schematic figure of the method of images showing the translocation of particle through a channel.

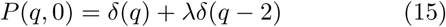

where *λ* determines the strength of the mirror image source. Assuming the initial condition of particle capture into the channel as: *P* (*q,* 0) = *δ*(*q*), and the absorbing boundary condition once the particle has completely translocated into trans side of the channel: *P* (*q* = 1*,t*) = 0. From this, the result for first passage probability density function (FP-PdF) [32] can be written as:

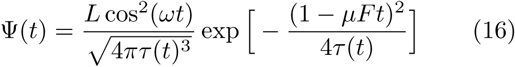

where Ψ(*t*) describes the probability density for the time that the particle spends in the channel. Figure (6) show the plot of the Eq. (16). The distribution of FPT displays many speaks at frequency *ω* = 3. Note that an increase in the frequency of oscillation *ω* causes an increase in the number of peaks in the distribution of first passage time (FPT). From this simple theory of particle translocation through a channel, we predict the peaks in the first passage time PdF when the transport process is perturbed by a simple time modulation of variance. Using the approximation of absorbing boundary condition at the trans side, *q* = *L*, the solution of Ψ(*t*) contains the trajectory of the random walker traveled to *q <* 0 side and come back to the region 0 < *q* < *L*. During particle translocation, there is a finite probability that the particle can retract into the cis side against the free energy barrier. Under such boundary condition the particle cannot enter the interval through an absorbing end point because its is instantly trapped through by the absorbing boundary.

**FIG. 6.**
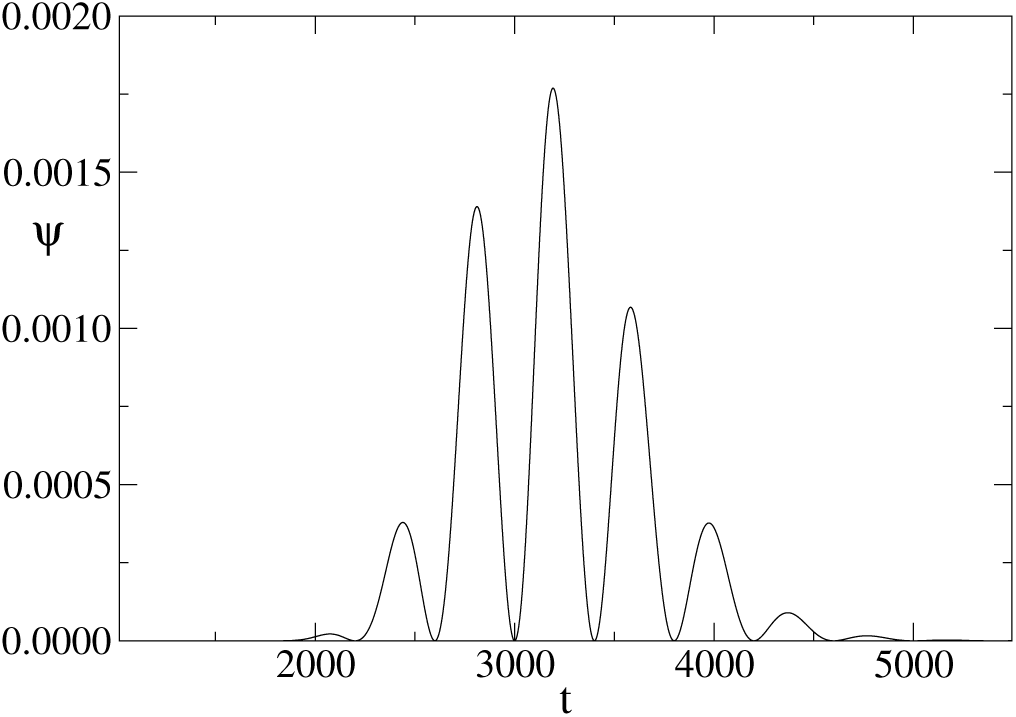
The distribution of first passage time computed via approximate solution of Langevin equation (Eq. (5)).

We anticipate that by introducing a simple time modulation in the variance of Langevin equation, the transport of single particle across a channel display many peaks in the distribution of translocation time. It is instructive to investigate the transport dynamics of dumbbell particle in order to understand the interplay of channel frequency and the oscillation of spring. In such toy model the dumbbell are described by harmonic oscillator, composed by two material points and connected via Hooke’s spring of elastic constant *k*.

Consider an elastic dumbbell particle made of two identical particle connected by a Hookean spring. The equation of motion for the particle 1 of a dumbbell is of the Langevin type

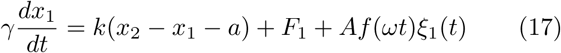

and similarly for particle 2

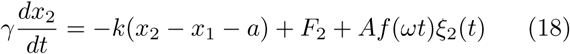

where *F*_1_ and *F*_2_ are constant forces acting on particle 1 and particle 2, respectively, and *k* is the Hook spring constant. Figure (7) illustrates the basic scheme of dumbbell particle translocation through a fluctuating channel.

**FIG. 7.**
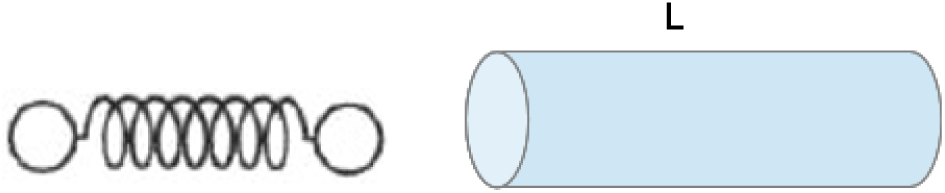
Two particle attached via Hooke’s spring translocating through a channel of length *L*.

Equations (17) and (18) are solved numerically using the standard Euler-Maruyama method [33]. Figure (8) describes the probability distribution function (PdF) of translocation time.

**FIG. 8.**
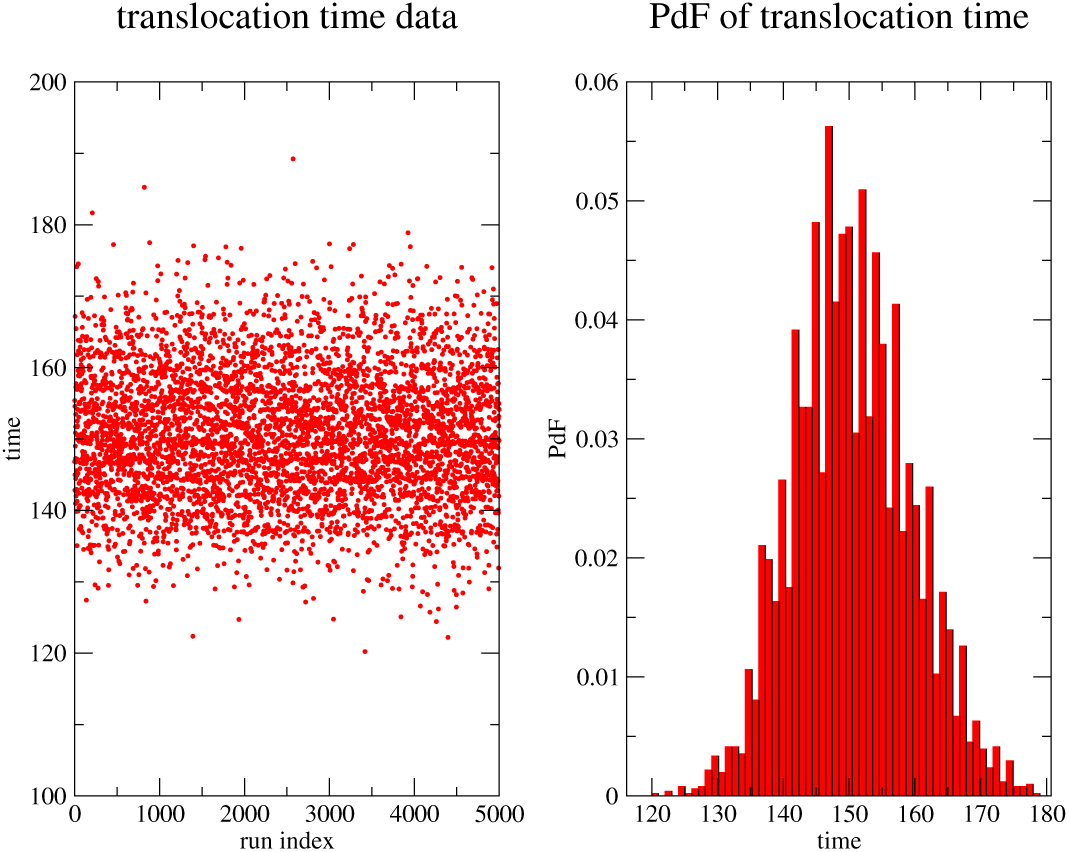
Right Panel: Probability distribution function of translocation time for two particle attached via Hooke’s spring *K* = 0.5 at frequency *ω* = 3. Left Panel: The dots represent the scatter data of translocation time.

In summary, we discuss a simple theoretical model to understand the protein translocation through a flickering pore. We study the dynamics of Brownian particle driven by a periodically driving force acting along the axis of the nanopore. Using Langevin equation with constants drift and time dependent variance, and first passage time theory we obtained the formalism for translocation time probability density function (PdF).

The author would like to thank U. M. B. Marconi, A. Vulpian and F. Cecconi for helpful discussion, and Institute of Complex Systems, National Research council of Rome for providing access to computational resources.

